# Broadband gamma-band EEG changes during magnetophosphene perception induced by 20 Hz magnetic field stimulation

**DOI:** 10.64898/2026.04.15.718626

**Authors:** Maëlys Moulin, Eléonore Fresnel, Julien Modolo, Nicolas Bouisset, Sofiane Ramdani

**Author notes:** Author to whom any correspondence should be addressed. **E-mail:**.

## Abstract

**Objective:** Magnetophosphenes are visual percepts induced by extremely low-frequency magnetic fields (ELF-MF; <300 Hz), yet the EEG expression of suprathreshold magnetophosphene-inducing stimulation remains poorly characterized and is not reliably captured by classical low-frequency markers. We tested whether suprathreshold 20 Hz tAMS is accompanied by broadband high-frequency EEG changes rather than focal oscillatory effects.

**Approach:** EEG was recorded in N=13 healthy volunteers during 20 Hz sinusoidal magnetic-field exposure delivered using transcranial alternating magnetic stimulation (tAMS) in a global-head configuration. Three conditions were analyzed: no exposure (0 mT), subthreshold (5 mT), and suprathreshold (50 mT). Gamma-band activity (30–80 Hz) was quantified using complementary spectral approaches, including aperiodic-adjusted gamma power.

**Main results:** Perception reports sharply dissociated the three conditions, with frequent perception at 50 mT only. Suprathreshold stimulation was associated with spatially distributed increases in gamma-band activity over frontal and occipito-parietal electrodes.

These effects persisted after aperiodic correction using two independent parameterization methods and did not exhibit a consistent narrowband peak, indicating broadband high-frequency changes. No focal power change over primary occipital electrodes remained significant after Bonferroni correction.

**Significance:** Suprathreshold magnetophosphene-inducing stimulation is not reliably captured by focal low-frequency EEG markers but is accompanied by distributed broadband high-frequency activity. Because stimulation intensity and perceptual reports were strongly coupled, these effects should be interpreted as EEG correlates of a suprathreshold stimulation-perception state rather than as isolated markers of perception.

## 1 Introduction

Magnetophosphenes and electrophosphenes denote visual experiences that arise without any optical input, triggered by time-varying magnetic or electric fields delivered to the eye and/or head [1, 2]. First described by d’Arsonval [3], these percepts have been extensively characterized in humans, with maximal sensitivity around 20 Hz [4, 5]. Subsequent work has examined their dependence on stimulation parameters, ambient conditions, and potential interaction sites along the visual pathway [6, 7].

Within the context of exposure to extremely low-frequency magnetic fields (ELF-MF; <300 Hz), magnetophosphenes have primarily been studied as a psychophysical threshold phenomenon [4, 6]. These data underpin threshold–frequency curves used in exposure assessment and contribute to the definition of safety limits [8–10]. In this framework, phosphene reports are typically reduced to binary perceptual judgments (*perceived* vs. *not perceived*) [2, 4, 11]. Similar threshold-based approaches are used in transcranial magnetic stimulation (TMS), where phosphene thresholds serve as operational markers of visual system excitability [12–14], and in transcranial alternating current stimulation (tACS), where phosphene thresholds have been mapped across frequencies and montages and contribute to phosphene-based safety considerations [10, 15, 16].

A key complication is that phosphenes may arise from different interaction sites (retinal vs. cortical). Empirical dissociation remains difficult across transcranial electrical stimulation paradigms [17, 18], although modeling supports a predominantly retinal origin under typical conditions [19]. For ELF-MF exposure, recent dosimetry and montage comparisons similarly support a predominantly retinal interaction site under global-head stimulation, making a sufficient direct cortical origin unlikely [2, 11].

In the context of non-invasive brain stimulation, phosphenes are not merely perceptual curiosities but stimulation-induced sensory phenomena that constrain tolerability, safety, and the interpretation of neuromodulatory effects. Despite well-characterized behavioral thresholds, however, their electrophysiological expression remains poorly characterized, particularly for magnetic-field exposures in the ELF range. EEG studies have not identified a robust focal occipital signature: subthreshold exposures show little or no reliable modulation of occipital alpha power, whereas studies targeting conscious magnetophosphene perception report changes at the level of large-scale functional organization, such as increased alpha-band integration, rather than local power shifts [20–22].

Determining whether suprathreshold tAMS produces focal sensory-like EEG responses or distributed broadband changes is therefore relevant for both mechanistic interpretation and stimulation safety.

Classical accounts of visual perception often relate perceptual reports to occipital alpha activity (8–12 Hz), interpreted as indexing fluctuations in cortical excitability and sensory gain [23–25]. Reduced prestimulus alpha power has been associated with improved visual detection and trial-to-trial perceptual variability [26–31]. By extension, one might expect magnetophosphene perception to be accompanied by changes in occipital alpha power.

However, this expectation is not supported empirically and is conceptually limited. Alpha–perception relationships depend strongly on task and stimulus properties and do not constitute a context-independent marker [29, 30, 32, 33]. Consistently, magnetophosphene studies do not report a reliable focal occipital alpha signature [20–22]. Moreover, evidence from TMS-induced phosphenes indicates that perceptual reports are associated with distributed and delayed responses rather than early, localized occipital activity [34–37]. Together, these findings suggest that phosphene perception is unlikely to be captured by a simple local oscillatory marker.

More broadly, current accounts of perception emphasize distributed and recurrent processing across cortical networks rather than isolated changes in a single sensory region [38–41]. In this framework, EEG signatures of perceptual access are expected to reflect large-scale and multiscale dynamics rather than focal low-frequency power modulations [42, 43]. This perspective is consistent with magnetophosphene studies reporting changes in large-scale functional organization rather than local spectral effects [22].

Magnetophosphene perception further differs from classical visual paradigms in that it arises without structured visual input. Under ELF-MF exposure, induced electric fields primarily affect the retina and lack retinotopic spatial organization [2, 11, 44]. Consequently, cortical activity is unlikely to be constrained by the same feedforward sensory drive as in photon-driven vision, and EEG signatures are not expected to replicate those observed in classical detection paradigms [45–50].

These considerations motivate the search for alternative EEG markers that go beyond local low-frequency oscillations. In particular, higher-frequency activity in the gamma range has been linked to population-level processing and large-scale integration, often manifesting as broadband rather than narrowband spectral changes [51–53]. In phosphene-related paradigms, such activity has been reported as distributed broadband patterns [54, 55]. Importantly, standard spectral measures conflate periodic and aperiodic components, and recent approaches allow these contributions to be separated [56, 57].

Here, we investigate EEG correlates of magnetophosphene perception by moving beyond classical alpha-based markers. We test whether perceptual reports are associated with broadband gamma-band activity (30–80 Hz) and whether these effects persist after separating periodic and aperiodic components. Rather than inferring a specific neural mechanism, our goal is to determine whether higher-frequency spectral features provide a more informative description of perception-related EEG changes in this context. We hypothesize that suprathreshold stimulation is associated with broadband gamma-band differences that remain detectable after aperiodic adjustment.

## 2 Materials and Methods

### 2.1 Participants

Fourteen healthy volunteers (7 females; mean age 25.8*±*5.27 years) participated in the study. An *a priori* sample size estimation was performed using G*Power [58] under a repeated-measures ANOVA approximation (three within-subject conditions, one group). Assuming a medium-to-large standardized effect size (*f* = 0.40), *α* = 0.05, and 80% power, this analysis yielded a minimum required sample size of 12 participants. To account for an anticipated attrition or exclusion rate of approximately 10% due to EEG data quality issues, 14 participants were therefore recruited. One participant was excluded from group-level analyses based on the predefined EEG quality-control procedure described below (Data quality control and subject exclusion; see Supplementary Figure 5), resulting in 13 participants included in the final analyses.

Since the primary analyses were conducted at the trial level using linear mixed-effects models and involved stringent multiple-comparison correction across electrodes, we also conducted a simulation-based sensitivity analysis using the simr package [59] in R (version 2026.01.0). Simulations preserved the trial-level structure (repeated trials nested within subjects, random intercept for subject) and applied an electrode-wise Bonferroni correction (*α* = 0.05*/*61). Under this conservative threshold, effects comparable to the smallest coefficients observed in the dataset were underpowered; however, assuming a moderate effect size on the transformed scale (e.g., *β* = 0.05 for the intensity contrast), a sample size of 13 participants provided approximately 80–90% power.

### 2.2 Experimental protocol

EEG data were acquired during experimental sessions using the stimulation system and protocol described by Legros et al. [2] and Fresnel et al. [11]. Following their terminology, we refer to the present 20 Hz global-head ELF-MF exposure (5-s sinusoidal magnetic-field) as transcranial alternating magnetic stimulation (tAMS). The technical details on the apparatus, coil geometry, field generation, dosimetry, safety architecture, and experimental control are reported in [11] and are not repeated here.

The present study focuses exclusively on EEG recordings obtained during the 20 Hz global-head condition. Each participant completed 55 trials corresponding to 11 magnetic-field flux density values ranging from 0 to 50 mT in 5 mT steps, each repeated five times and pseudo-randomized across trials. Magnetic-field amplitudes are RMS values. For readability, we denote intensities in mT throughout the manuscript (RMS implied unless stated otherwise). After each exposure, participants provided a behavioral report of magnetophosphene perception via a button press (binary response: perceived vs. not perceived). Participants were seated comfortably in complete darkness with eyes closed throughout the experiment (Figure 1). Consecutive stimulation trials were separated by a 5 s interstimulus interval. This interval was chosen in the original psychophysical protocol to reduce immediate carry-over effects while maintaining participant vigilance and limiting the overall duration of the session [2, 11]. Stimulation intensities were pseudo-randomized across trials, which reduced the likelihood that any residual effect of a given intensity would systematically bias a specific subsequent condition. However, no dedicated washout manipulation was included in the protocol.

**Figure 1.**
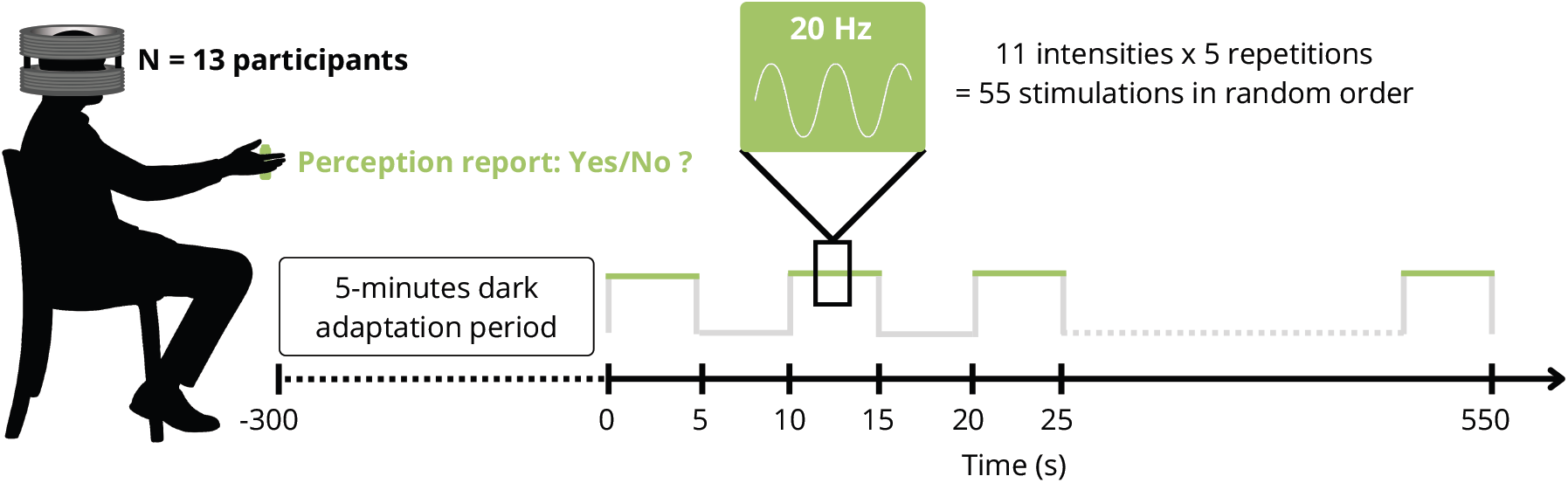
Overview of the experimental protocol. After a 5-min dark-adaptation period, participants (N = 13) received 55 sinusoidal ELF-MF stimulations at 20 Hz (11 intensities*×* 5 repetitions) presented in pseudo-random order. Each stimulation lasted 5 s and was followed by a binary perceptual report (Yes/No) with a button press.

For the EEG analyses reported here, we focused only on three conditions: a no-field control (0 mT), a subthreshold exposure (5 mT), and a suprathreshold exposure (50 mT). These conditions were selected to clearly contrast non-perceptual and perceptual states while minimizing inter-individual heterogeneity near threshold. Intermediate intensities were not included in the primary EEG analyses because they sampled the psychophysical transition region and therefore produced heterogeneous perceptual states across participants. With only five repetitions per intensity, these intermediate levels did not provide sufficient trial numbers to reliably separate perceived from non-perceived trials within each intensity and subject. The present manuscript therefore focused on three predefined conditions providing clearly interpretable stimulation-perception states: no-field control (0 mT), weak subthreshold exposure (5 mT), and robust suprathreshold exposure (50 mT). A full dose-response analysis of the intermediate intensities will require dedicated modeling of individual perceptual thresholds and is beyond the scope of the present report. Behavioral results (button-press reports) are reported in the Results section. No modification of the stimulation hardware, experimental timing, or procedure relevant to EEG acquisition was introduced relative to the protocol described in [11].

### 2.3 EEG recording

EEG recordings were acquired at a sampling rate of 10 kHz using a 64-channel MRI-compatible Neuroscan system (Compumedics Supplies, Charlotte, NC, USA) arranged according to the international 10–10 system. The high sampling rate was used to accurately capture the rapid transients associated with the magnetic field stimulation artifact and to enable precise detection of stimulation onset times prior to subsequent downsampling. Signals were recorded with CPz as the reference electrode and AFz as ground. Additional bipolar channels were used to monitor ocular (EOG) and cardiac (ECG) activity for online monitoring and offline quality control. Online artifact correction options implemented in the Neuroscan acquisition software were enabled, including blink noise reduction and ocular artifact reduction based on bipolar EOG channels (VEOG/HEOG). The spatial SVD filter was explicitly disabled during acquisition.

### 2.4 EEG preprocessing

Continuous EEG data were imported into MATLAB^®^ (MathWorks Inc., Natick, MA, USA) using the EEGLAB toolbox. Data were re-referenced offline to the average of the left and right mastoids (M1/M2). For each participant, a detection channel was selected as the channel with the largest stimulation artifact (priority was given to a dedicated MF channel), using an automatic heuristic with manual override in a few problematic cases. Stimulation onsets were then detected on this raw detection channel using an envelope-based detector. Based on the detected onset indices and the individual randomization list of stimulation intensities (55 trials per subject, including 0–50 mT ), we reconstructed 5-s epochs for all EEG channels. Epochs corresponding to real stimulations (intensity ≠0 mT) were time-locked to MF exposure onset. For the control condition (0 mT), epochs were constructed using control time windows, defined as 5-s segments selected within*±*10 s of a real stimulation onset but outside any field exposure period. Control windows were required to be free of stimulation onset markers and did not overlap with any real stimulation epoch, ensuring comparable temporal context across conditions.

Each 5-s epoch was multiplied by a Hann window and band-pass filtered between 30 and 80 Hz using a zero-phase finite-impulse response filter. To minimize edge artifacts, only the central 1–4 s of the filtered epoch was retained for spectral analysis. Residual narrowband components at 40, 50, and 60 Hz were removed by applying a cascade of second-order IIR notch filters (zero-phase `filtfilt` implementation) with an effective half-width of approximately*±*1.5 Hz around each center frequency. Frequency bins located within *±*1.5 Hz of 40, 50, and 60 Hz were excluded to minimize residual stimulation- and power-line–related artifacts while preserving broadband gamma activity outside these narrow contamination regions. Downsampling was applied to reduce computational cost while preserving the frequency range of interest (30–80 Hz) given the post-filtered signal bandwidth. The notch-cleaned signal was then downsampled using the `resample` function by a factor of 5 (with the effective sampling frequency adjusted accordingly: *F*_s,eff_ = *F*_s_*/*5 = 2 *kHz*).

Independent Component Analysis (ICA) was not applied since our analyses focused on the 30–80 Hz band on short stimulation-locked epochs after online ocular artifact reduction and notch filtering. Remaining artifacts were addressed through manual epoch selection and subject-level quality control. For statistical analyses, only the epochs corresponding to 0 mT (control), 5 mT, and 50 mT conditions were retained, resulting in 15 epochs of 3-s duration (1–4 s) per electrode per subject. Three electrodes (C3, Pz, and PO8) were excluded from electrode-wise statistical analyses because they repeatedly exhibited poor signal quality across several participants (high impedance).

### 2.5 Frequency analysis

Power spectral densities (PSDs) were estimated with Welch’s method [60] in MATLAB^®^, using 1-s Hann windows with 50% overlap, computed on the downsampled signal. Analyses focused on the 30–80 Hz gamma band. Because the signal was band-pass filtered to the gamma range prior to spectral estimation, we did not compute across-band relative power measures. Instead, we quantified absolute gamma power as the integral of the PSD between 30 and 80 Hz. Complementary aperiodic parameters and aperiodic-adjusted gamma power were extracted as described below.

### 2.6 Aperiodic–periodic spectral decomposition

An aperiodic background was parameterized using the *specparam* approach (formerly FOOOF) [56], implemented via a dedicated Python environment called from MATLAB^®^. Peak width limits were set to 1–12 Hz, with a maximum of six peaks and a peak detection threshold of two standard deviations above the aperiodic fit. For each trial, electrode, and stimulation intensity, *specparam* returned the aperiodic exponent and offset, as well as gamma peak center frequency and amplitude when detected.

*Specparam* goodness-of-fit (*R*^2^) was extracted to characterize model performance across conditions and is reported for quality-control purposes, but was not used as an exclusion criterion. *R*^2^ values were examined to assess whether fit quality differed systematically across stimulation conditions (Supplementary Figure 6). Because *specparam* models the spectrum as an aperiodic component plus putative Gaussian-like periodic peaks, goodness-of-fit values were expected to be limited when gamma-band activity was broadband and did not contain a stable narrowband peak. Moreover, spectra were parameterized within a restricted 30–80 Hz window after exclusion of narrow frequency bands around 40, 50, and 60 Hz. Therefore, *specparam* goodness-of-fit values were interpreted as local quality-control descriptors rather than as evidence for a complete model of the full power spectrum.

To avoid relying on this peak-based parameterization alone, we also estimated the aperiodic component using *log–log* linear regression and subtracted it from the PSD to obtain a residual spectrum [61]. This complementary approach provided an assumption-light estimate of aperiodic-adjusted gamma power that did not rely on explicit oscillatory peak modeling. Using both the *specparam* and *log–log* approaches enabled us to assess whether gamma-band effects were robust to different aperiodic-correction strategies.

All reported gamma-band metrics included raw gamma power and two estimates of aperiodic-adjusted gamma power, derived respectively from the *specparam* and *log–log* approaches and defined as positive residual power above the local aperiodic fit within the 30–80 Hz range, excluding frequency bins within*±* 1.5 Hz of 40, 50, and 60 Hz. Aperiodic parameters were reported descriptively, together with gamma peak center frequency when detected and *specparam* goodness-of-fit (*R*^2^). All metrics were derived from the same notch-cleaned, downsampled epoch segments.

### 2.7 Data quality control and subject exclusion

EEG data quality was evaluated at the level of whole recordings rather than by dropping individual channels. In practice, most participants had a few noisy electrodes, but affected channels differed from one subject to another, which made any fixed channel-rejection rule arbitrary. More importantly, once the preprocessing pipeline had been applied, channels of poor quality in the raw data often produced spectral and dynamical estimates comparable to those obtained from visually clean channels in other participants. Relying on raw visual appearance alone therefore did not provide a stable or reproducible criterion for exclusion (Figure 2).

**Figure 2.**
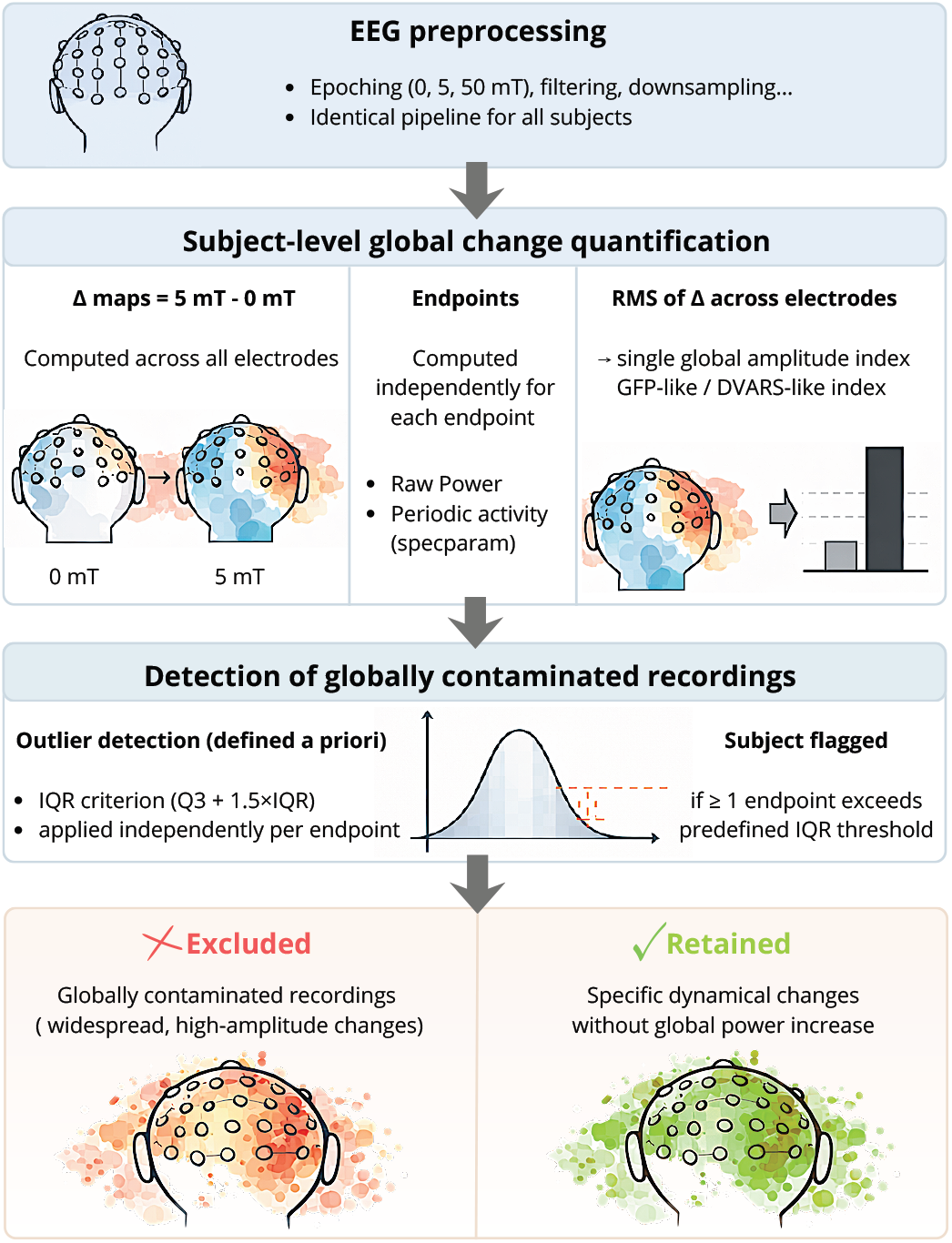
Overview of the subject-level EEG quality-control and exclusion workflow. EEG signals were first preprocessed using an identical pipeline for all participants, including epoching (0, 5, and 50 mT conditions), filtering, and downsampling. For each subject, condition-related difference maps (Δ = 5 mT− 0 mT ) were computed across all electrodes. Global change magnitude was then quantified independently for multiple EEG outcome measures, including raw gamma power and aperiodic-adjusted gamma power estimated using *specparam*. For each EEG outcome measure, the root mean square (RMS) of the Δ values across electrodes was computed, yielding a single subject-level index of global amplitude change (conceptually analogous to GFP- or DVARS-like metrics). Subjects were flagged using an *a priori*–defined outlier detection criterion based on the interquartile range (IQR; threshold = *Q*_3_ + 1.5 *×* IQR), applied independently to each EEG outcome measure. A recording was classified as globally contaminated if at least one EEG outcome measure exceeded its predefined threshold. This procedure was designed to selectively identify recordings dominated by widespread, high-amplitude changes consistent with global artifacts, while retaining subjects exhibiting spatially structured and physiologically plausible dynamical modulations without a global power increase.

Given the constraints specific to ELF-MF recordings, we opted for a subject-level quality-control index that targets globally abnormal responses rather than local channel noise. For each participant, we computed stimulation-related difference maps (Δ = 5 mT − 0 mT) across all electrodes for both spectral EEG outcome measures derived from PSD analyses. The root mean square (RMS) of each Δ map was then used as a compact summary of how large and spatially widespread the stimulation-related change was for that subject. This RMS index plays a role similar to global-field-power–like summaries in EEG [62, 63] and is conceptually close to DVARS-type measures used to flag global deviations in fMRI [64, 65]. The rationale was to identify recordings dominated by large, spatially diffuse changes that are more consistent with contamination than with plausible neurophysiological effects.

For each EEG outcome measure (raw gamma power and aperiodic-adjusted gamma power estimated using the *specparam* approach), subjects with unusually large RMS values were flagged using an interquartile-range rule (*Q*_3_ + 1.5*×*IQR). A stricter cutoff (*Q*_3_ + 3*×*IQR) was checked as a sensitivity analysis and led to the same exclusion decision. Only extreme high-RMS values were considered problematic, because low RMS simply reflects weak or absent global effects rather than contamination.

RMS values were also *z*-scored within each EEG outcome measure to compare how strongly individual subjects contributed across metrics. These *z*-scores were used descriptively to visualize cross-EEG outcome measure consistency and were not used as formal exclusion thresholds. For power-based metrics, RMS was computed on fourth-root–transformed values to remain consistent with the statistical models. All quality-control criteria were defined *a priori* and applied blind to subject identity and group-level outcomes. Subjects flagged by this procedure were removed from subsequent analyses (see Supplementary Figure 5).

### 2.8 Statistical analysis

All analyses were performed in R (R Foundation for Statistical Computing, Vienna, Austria). Three within-subject contrasts were evaluated: 0 vs. 5 mT, 0 vs. 50 mT, and 5 vs. 50 mT. The 50 mT vs. 0 mT contrast was considered the primary comparison for characterizing suprathreshold stimulation effects. The 5 mT vs. 0 mT contrast served to verify that weak subthreshold exposure did not induce detectable gamma-band changes, whereas the 50 mT vs. 5 mT contrast was retained as a secondary control comparison to assess whether suprathreshold effects persisted relative to a nonzero but subthreshold field condition. Since gamma-band power metrics exhibit skewness typical of chi-squared–distributed quantities, all power-based measures were transformed using the fourth root, a transformation known to provide an excellent approximation to normality across degrees of freedom [66].

For each gamma-band metric, each electrode, and each contrast, the effect of stimulation intensity was tested using linear mixed-effects models (lme4 package), with intensity as a fixed factor (two levels) and a random intercept per subject. The *p*-value was extracted for the intensity effect, and a Bonferroni correction was applied, resulting in an electrode-wise significance threshold of *α* = 0.05*/*61 (61 electrodes), i.e., *p≈*8.2*×*10^−4^. In parallel, a paired Cohen’s *d*_*z*_ effect size was computed from within-subject differences.

Electrode-wise mean differences (difference between intensities), Bonferroni-corrected *p*-values, and direction *×*significance indices were visualized as interpolated scalp topographies (`eegUtils` package).

Complementary frequency-wise analyses were also performed at the ROI level. For each ROI (parieto-occipital and frontal), PSD values were averaged across the electrodes defining the ROI for each subject, condition, and frequency bin. Linear mixed-effects models identical to those described above were fitted independently at each frequency bin in the 30–80 Hz range. These analyses focused on the 50 mT vs. 0 mT contrast. Resulting *p*-values were Bonferroni-corrected across frequency bins within each ROI and spectral parameterization.

## 3 Results

### 3.1 Behavioral results: intensity-dependent perception

Magnetophosphene perception strongly depended on stimulation intensity. No perception was reported at 0 mT (0/65 trials), and only a single trial resulted in perception at 5 mT (1/65 trials; 1.5%), whereas perception was frequent at 50 mT (57/65 trials; 87.7%). This sharp behavioral dissociation justified contrasting the control (0 mT), subthreshold (5 mT), and suprathreshold (50 mT) conditions in subsequent EEG analyses. When reported, magnetophosphenes were described as brief, flickering visual sensations occurring in the absence of light. Participants typically reported diffuse, poorly delimited percepts rather than stable shapes or spatially structured patterns. The percepts were described as colorless and transient, and their temporal occurrence was aligned with the stimulation periods. Several participants spontaneously reported that the sensations appeared to involve peripheral vision; however, no formal spatial rating was collected.

### 3.2 Spectral profiles of gamma-band activity

Gamma-band activity was elevated in the suprathreshold condition (50 mT) relative to the control (0 mT) and subthreshold (5 mT) conditions in both parieto-occipital (O1, O2, Oz, PO1–PO6, POz, P1–P6 and Pz) and frontal (FP1, FP2, AF3, AF4, F1–F4 and Fz) ROIs (Figure 3). This elevation extended over a broader portion of the gamma range in frontal electrodes than in occipital ones, although the overall spectral profiles remained comparable across regions. The figure displays raw spectra as well as spectra corrected for the aperiodic component using both the *specparam* approach and the *log–log* approach.

**Figure 3.**
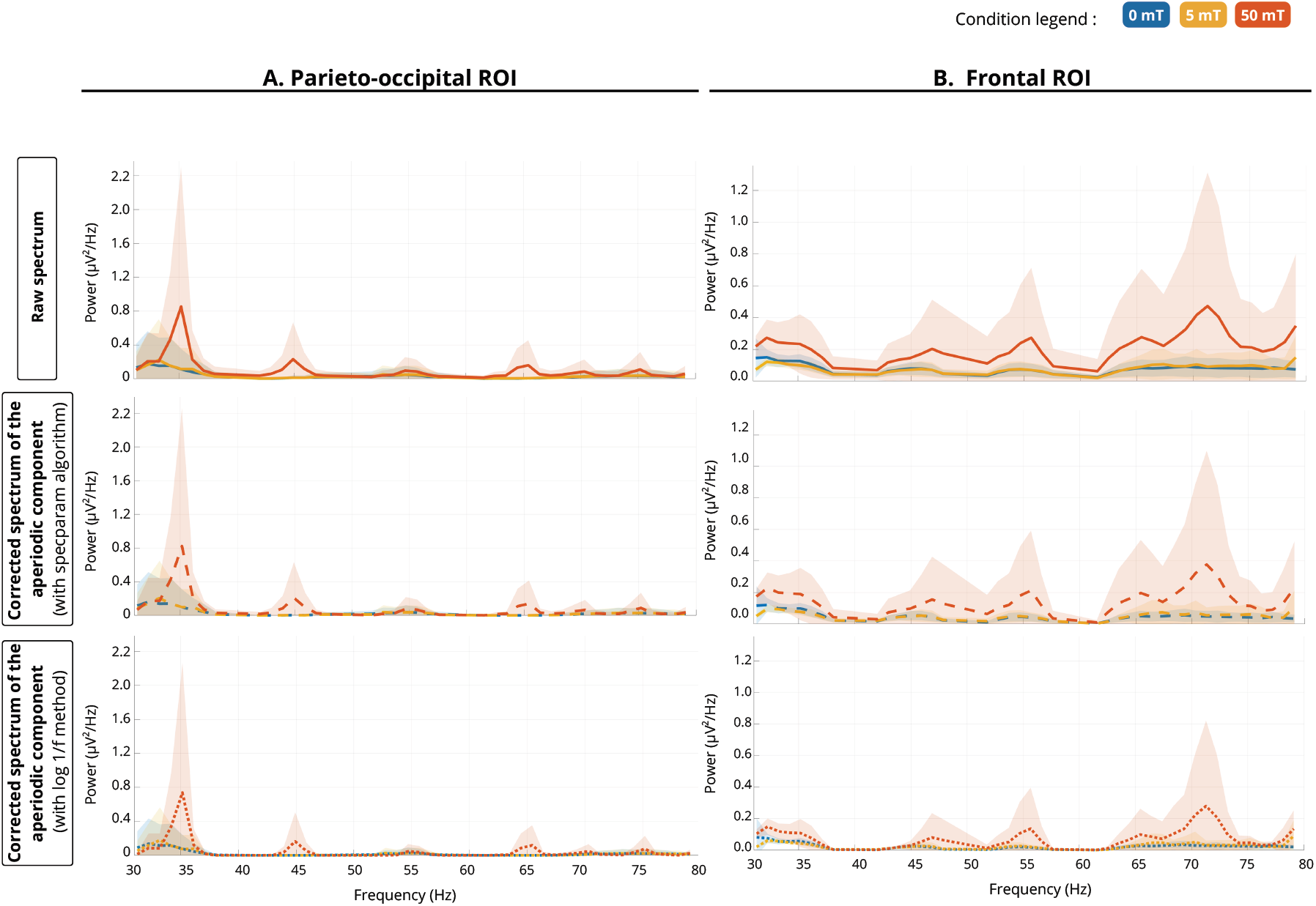
PSD in parieto-occipital (A) and frontal (B) regions of interest. Group-averaged PSD estimates are shown for parieto-occipital (left) and frontal (right) ROIs in the 30–80 Hz range. Colors denote stimulation intensity: 0 mT (blue), 5 mT (yellow), and 50 mT (red). Line styles indicate spectral representations: solid lines correspond to raw power spectra; dashed lines show aperiodic-adjusted gamma power estimated using the *specparam* approach; dotted lines show aperiodic-adjusted gamma power estimated after removal of the aperiodic background using the *log–log* approach. Shaded areas indicate the 95% CI across subjects. Across conditions, spectra show a broadband elevation of high-frequency power at 50 mT, more pronounced in frontal than in occipital electrodes, whereas modulation of the periodic component remains more limited and spatially heterogeneous.

To quantify these spectral differences, frequency-wise mixed-effects analyses comparing the 50 mT and 0 mT conditions were performed across the 30–80 Hz range.

Effects in the parieto-occipital ROI were relatively restricted, with localized differences between the 50 mT and 0 mT conditions observed around 34–36 Hz (Table 1). By contrast, the frontal ROI exhibited more consistent and spectrally broader differences across parameterizations, including a peak in the lower gamma band (approximately 32–37 Hz) and additional peaks at higher frequencies, around 68–74 Hz (Table 1).

**Table 1.**
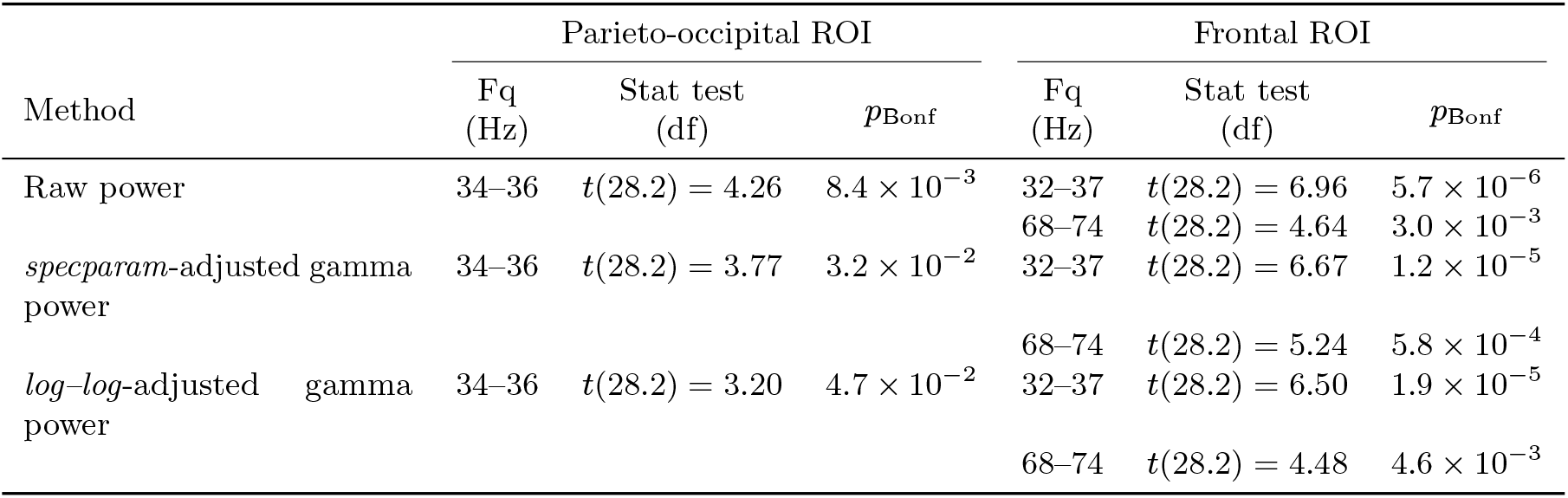
Peak gamma-band effects for the comparison between the 50 mT and 0 mT conditions across spectral parameterizations. Frequencies (Fq) indicate the bin showing the strongest statistical effect within each ROI. Reported *p*_Bonf_ values are Bonferroni-corrected across frequency bins within each ROI and spectral parameterization.

Decomposing spectra into periodic and aperiodic components indicated that the suprathreshold effect was not reducible to a uniform shift of the spectral background. After correcting for the aperiodic (1/f-like) component, gamma-band differences between 50 mT and the control condition (0 mT) remained visible in the corrected spectra, although attenuated. At 50 mT, aperiodic correction removed a substantial proportion of broadband gamma power: on average 43.2% in the frontal ROI and 41.2% in the occipital ROI using the *specparam* approach, compared with about 65.6% in both ROIs with the *log–log* approach.

Consistent with this pattern, no stable narrowband gamma peak emerged across subjects or regions: instead, an elevation of high-frequency content rather than the appearance of a specific oscillatory component was observed.

### 3.3 Spatial distribution of gamma-band effects

Electrode-wise contrasts (Figure 4) confirmed that 5 mT did not produce spatially consistent changes relative to control across any spectral representation. No electrode survived Bonferroni correction for the 5 mT vs. 0 mT contrast in any of the three gamma estimates.

**Figure 4.**
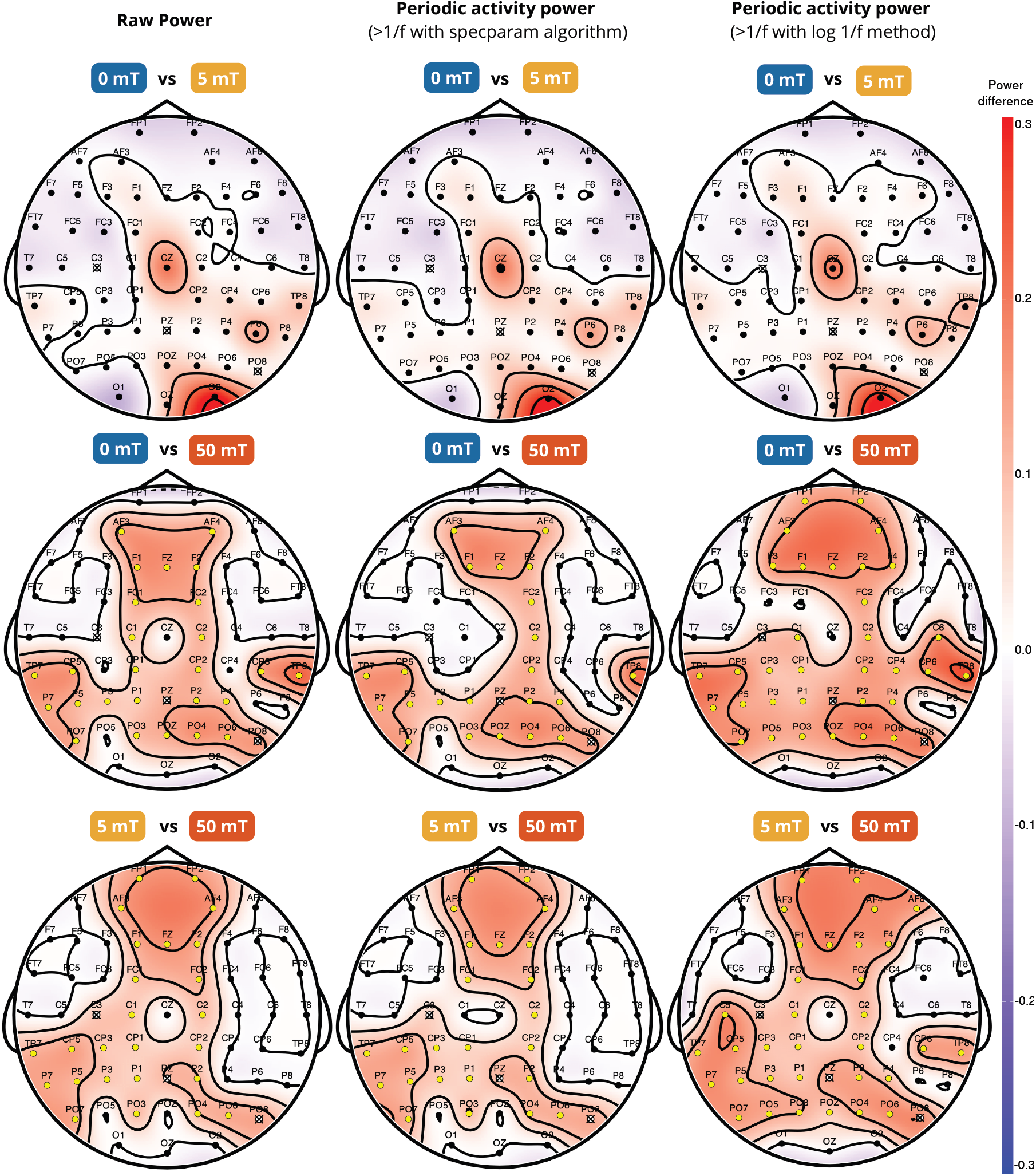
Scalp topographies of gamma-band power differences across stimulation intensities and spectral representations. Rows show within-subject contrasts (5 mT vs. 0 mT, 50 mT vs. 0 mT, and 50 mT vs. 5 mT), and columns correspond to raw gamma power (left), *specparam*-adjusted gamma power (middle), and log–log adjusted gamma power (right). Color represents the difference between conditions (Δ, higher− lower; fourth-root transformed scale). Yellow markers indicate electrodes remaining significant after Bonferroni correction, black markers indicate electrodes not surviving correction, and white markers with a black cross indicate electrodes excluded from statistical analyses (C3, Pz, and PO8) and shown for topographical completeness only. Full electrode-wise statistical results (*p*_Bonf_ and *d*_*z*_) are provided in Supplementary Figure 7.

In contrast, 50 mT was associated with widespread increases in raw gamma power, with a clear centro-frontal predominance. After aperiodic adjustment, the overall pattern remained present but became less spatially extensive and more heterogeneous, and the exact distribution depended on the parameterization method used to estimate periodic activity (*specparam* vs. *log–log* subtraction).

The 50 mT vs. 5 mT contrast was retained as a control comparison to test whether the suprathreshold effect was attributable merely to the presence of a nonzero magnetic field. As expected from the absence of significant differences between 5 mT and 0 mT, the 50 mT vs. 5 mT contrast closely resembled the 50 mT vs. 0 mT contrast. This similarity was therefore not interpreted as independent evidence for a distinct effect, but rather as a consistency check showing that weak subthreshold exposure did not produce a scaled-down version of the suprathreshold response. The 50 mT vs. 0 mT contrast remains the primary comparison for characterizing suprathreshold stimulation effects.

Overall, the strongest effects were observed over fronto-central and parieto-occipital regions, whereas the primary occipital electrodes O1, O2, and Oz did not survive multiple-comparison correction in any representation. For the 50 mT vs. 0 mT contrast, all Bonferroni-corrected values remained non-significant at O1, O2, and Oz across raw and aperiodic-adjusted representations (*p*_Bonf_*≥*0.05), despite moderate-to-large effect sizes (*d*_*z*_ ∈ [0.28, 0.88]) that did not reach significance after correction, suggesting substantial variability across participants. For the 50 mT vs. 5 mT contrast, corrected values at O1, O2, and Oz were likewise non-significant across representations (*p*_Bonf_ ≥0.06), with effect sizes ranging from negligible to moderate (*d*_*z*_ ∈ [−0.01, 0.29]).

For completeness, full electrode-wise statistical results (Bonferroni-corrected *p*-values and paired effect sizes) are provided in Supplementary Figure 7.

## 4 Discussion

The present study highlights a scalp-level electrophysiological signature accompanying suprathreshold magnetophosphene-inducing stimulation. This stimulation-perception state was associated with broadband high-frequency EEG increases that were spatially distributed and persisted after aperiodic adjustment. These findings extend prior work showing that magnetophosphene perception is not reliably captured by focal low-frequency markers [20–22].

### 4.1 Broadband high-frequency activity as the primary EEG correlate

The observed gamma-range effects are not consistent with a canonical oscillatory response. They are broadband, lack a reproducible peak across subjects and regions, and remain detectable after aperiodic adjustment. This pattern is therefore better described as an increase in high-frequency content rather than the recruitment of a specific gamma rhythm, consistent with previous work showing that much EEG “gamma” reflects non-narrowband activity [51, 52]. Broadband high-frequency activity has also been linked to aggregate population firing and synaptic activity, rather than to oscillatory synchronization per se [51]. Importantly, the effect is not reducible to a simple upward shift of the spectral background. Although aperiodic parameters estimated within a restricted 30–80 Hz window should be interpreted cautiously as local descriptors rather than global 1/f properties [56, 57], suprathreshold differences remained detectable after aperiodic correction using two independent approaches. The most conservative interpretation is therefore that, within the analyzed band, the effect cannot be explained by uniform amplitude scaling alone.

A critical limitation concerns the interpretation of high-frequency activity. EEG gamma-band signals are particularly vulnerable to non-neural contamination, including craniofacial muscle activity or subtle movements [51]. In addition, the 50 mT condition necessarily confounds high stimulation intensity with high perception frequency, making it impossible to fully disentangle intensity-dependent and perception-related contributions in the present design. Nevertheless, several features argue against a purely nonspecific intensity-driven or artifactual explanation. If broadband gamma increases were driven only by the physical intensity of a relatively global stimulation, or by a uniform stimulation-related artifact, one would expect a more spatially homogeneous scalp distribution. Instead, the suprathreshold response was spatially structured, with stronger effects over fronto-central and parieto-occipital regions, no focal effect over primary occipital electrodes remained significant after Bonferroni correction, and attenuation after aperiodic adjustment. Moreover, weak subthreshold stimulation did not produce a scaled-down version of the 50 mT response, as no consistent effects were observed for the 5 mT vs. 0 mT contrast. These observations suggest that the broadband gamma increase cannot be reduced to the mere presence of a magnetic field, to a trivial global intensity effect, or to a generic broadband noise increase. However, residual non-neural contamination cannot be excluded, and future studies combining individualized thresholding, graded stimulation around threshold, and perceptual intensity ratings will be required to separate physical intensity effects from perceptual-state effects.

This interpretation motivates considering why such effects do not appear as a focal occipital marker.

### 4.2 Absence of a robust focal occipital marker

A central result of the present study is the absence of effects at O1, O2, and Oz remaining significant after Bonferroni correction, despite reliable perceptual reports and widespread high-frequency changes. This absence should not be interpreted as a null result in the sense of absent visual-system involvement, but rather as evidence that magnetophosphene-inducing stimulation does not produce a stable focal occipital power change detectable at the scalp level.

This finding is consistent with the phenomenology and likely origin of magnetophosphene perception. Magnetophosphenes are typically diffuse, peripheral, and not driven by structured visual input [1, 2, 11]. Under ELF-MF exposure, converging dosimetric and psychophysical evidence supports a predominantly retinal interaction site, with limited contribution from direct occipital stimulation [2, 11, 67]. Neural activity reaching the cortex may therefore already be spatially and temporally diffuse, reducing the likelihood of a clear retinotopic scalp pattern.

This interpretation also aligns with previous ELF-MF EEG studies, which have not identified a reproducible focal occipital spectral marker of magnetophosphene perception [20–22]. In an acute 60 Hz EEG/fMRI study, no significant modulation of occipital alpha power and no detectable BOLD changes were reported under short exposures up to 7.6 mT, and no magnetophosphene perception was reported [20]. Conversely, when conscious magnetophosphene perception was explicitly targeted, it was associated with large-scale network reorganization, including increased alpha-band integration and altered beta-band segregation, rather than with a localized occipital spectral effect [22]. Together, these findings suggest that magnetophosphene-related EEG activity is more readily expressed at the level of distributed spectral or network-level changes than as a single focal occipital mechanism.

Earlier modeling work further supports the idea that ELF-MF effects may not manifest as simple local cortical responses. Biophysical models suggest that magnetic-field exposure may influence neural activity through distributed and multi-scale mechanisms, potentially involving indirect modulation of synaptic processes rather than direct localized activation [68]. This interpretation is also consistent with the limits of extrapolating from photon-driven visual paradigms: classical alpha-based gating mechanisms rely on structured, retinotopic input [23–25], whereas magnetophosphene perception arises without such feedforward constraints.

Importantly, the absence of a focal occipital signature should not be interpreted as evidence that early visual cortex is not involved in magnetophosphene perception. Rather, our results indicate that any potential involvement of early visual areas does not manifest, at the scalp level, as a stable and spatially focal gamma-band power change over primary occipital electrodes. This distinction is important because scalp EEG provides an indirect and spatially blurred measure of cortical activity, particularly for generators located within folded cortical structures such as the calcarine sulcus [69, 70]. In addition, simultaneous activation of opposite sulcal banks may lead to partial signal cancellation, and even in controlled retinotopic paradigms, identifying V1 generators from scalp EEG remains challenging [45, 71].

Taken together, these results suggest that magnetophosphene perception does not produce a robust focal occipital EEG marker at the scalp level, even if early visual areas are engaged. Instead, its electrophysiological expression appears to involve distributed dynamics that are only partially captured by classical focal spectral approaches.

### 4.3 Broadband activity as a large-scale state-dependent effect

The spatially distributed nature of the identified broadband gamma increase suggests that the observed effect reflects a state-dependent reconfiguration of ongoing activity rather than a localized sensory response. In this context, broadband high-frequency changes are better described as an increase in high-frequency/aperiodic activity without a stable spectral peak than as the recruitment of a task-specific gamma oscillator [51, 56, 72]. This spectral profile is more consistent with broader population-level activity than with a coherent rhythmic process at a fixed frequency [56, 61, 72].

Consistently, aperiodic spectral properties have been linked to excitation–inhibition balance and sensory variability [73], and gamma-range activity appears strongly shaped by stimulus and task context rather than reflecting a single invariant oscillatory mechanism [51, 52].

The frontal contribution to the observed effects is also compatible with a distributed, state-dependent interpretation. Frontal activity may reflect perceptual inference, report-related processes, or broader changes in arousal and expectancy associated with suprathreshold stimulation [39, 74]. More generally, internally generated percepts and false alarms can recruit neural signatures typically associated with conscious perception even in the absence of external sensory input [75, 76]. In the present paradigm, where perception emerges without patterned visual stimulation, magnetophosphene perception may therefore engage large-scale processes extending beyond early sensory encoding, consistent with theories of conscious access and prior magnetophosphene connectivity findings [22, 39, 40].

This interpretation remains cautious. High-frequency EEG signals are vulnerable to non-neural contamination, including craniofacial muscle activity and subtle movements [51, 72], and the present design cannot definitively separate neural from non-neural contributions. Nevertheless, the spatially structured pattern, its consistency across spectral representations, and its persistence after aperiodic adjustment argue against a purely generic broadband noise increase. The observed broadband gamma increase should therefore be interpreted as a correlate of the suprathreshold stimulation-perception state, rather than as a direct mechanistic signature of its neural origin.

### 4.4 Methodological constraints

Several aspects of the present design constrain the scope of interpretation. Analyses were restricted to the 30–80 Hz range and based on short (3 s) epochs, which improves sensitivity to high-frequency effects but precludes conclusions about cross-frequency interactions or global spectral structure.

Aperiodic parameters estimated within this window should therefore be interpreted as local descriptors, rather than global spectral properties [56, 57]. In addition, the binary contrast between clearly non-perceptual (0 and 5 mT) and perceptual suprathreshold (50 mT) conditions reduces ambiguity in classification but prevents inference about dose–response relationships. Future studies using individualized threshold-based sampling, more repetitions around threshold, and trial-by-trial perceptual intensity ratings will be required to determine whether broadband EEG changes scale with subjective perceptual strength. Finally, recordings acquired under magnetic-field exposure remain susceptible to residual non-neural contamination, particularly in the high-frequency range. Although several features of the results argue against a trivial artifact explanation, a definitive separation between neural and non-neural sources cannot be achieved with the present data.

The 5-s interstimulus interval represents an additional methodological constraint. Although this interval was used to reduce immediate carry-over effects and trial order was pseudo-randomized across stimulation intensities, the present design did not include a dedicated washout analysis or variable interstimulus intervals. Therefore, delayed or cumulative effects of stimulation cannot be fully excluded. Future studies could explicitly test carry-over effects by including longer or jittered interstimulus intervals and by modeling the influence of the preceding stimulation intensity on both perception reports and EEG activity.

### 4.5 Implications

These findings have two main implications. First, EEG markers classically associated with visual perception cannot be assumed to generalize to neuromodulation paradigms in which percepts arise without structured visual input. In magnetophosphene perception, the expectation of a focal occipital low-frequency signature appears both empirically fragile and conceptually mismatched to the sensory context [2, 11, 20–22]. Second, EEG analyses in this context may benefit from moving beyond classical band-power approaches toward descriptions of spectral structure and distributed state-dependent dynamics [56, 72]. Future work should extend this approach by incorporating temporal and nonlinear descriptors, such as entropy or recurrence-based analyses, to better characterize the global dynamics underlying perceptual states [77, 78].

### 4.6 Conclusion

Under 20 Hz ELF-MF exposure, magnetophosphene perception was associated with distributed broadband high-frequency EEG changes, while prior work has not identified a robust focal low-frequency marker over early visual cortex. Behavioral reports clearly dissociated non-perceptual conditions (0 and 5 mT) from the perceptual suprathreshold condition (50 mT). At the EEG level, suprathreshold stimulation was characterized by a spatially distributed increase in 30–80 Hz activity, without a consistent narrowband gamma peak and with no correction-surviving effects at primary occipital electrodes (O1, O2, Oz). Taken together, these findings indicate that suprathreshold magnetophosphene-inducing tAMS is not captured by classical focal spectral markers at the scalp level, but is instead accompanied by distributed broadband changes in ongoing EEG activity.

Importantly, the absence of correction-surviving effects at O1, O2, and Oz should not be interpreted as evidence against early visual cortex involvement, but rather as evidence that such involvement, if present, is not expressed as a stable focal power change detectable at the scalp level. This challenges standard assumptions derived from visual neuroscience and suggests that the neural expression of perception under neuromodulation may rely on large-scale, state-dependent dynamics rather than localized oscillatory mechanisms. Future work will determine whether these perception-related effects are also expressed in the temporal organization of EEG activity, using complementary dynamical descriptors such as entropy or recurrence-based measures.

## Acknowledgments

We gratefully acknowledge the late Dr. Alexandre Legros (†2025). His leadership, scientific vision, and scientific rigor were instrumental in shaping this work and continue to inspire our research.

We thank Simon Pla for his invaluable technical assistance throughout the study. We also thank the students who contributed their time and effort: Marvin Penault, Juliette Blaquart, Maorie Laporte, Tristan Dangoumau, and Hocine Chouaki.

## Funding

This work was supported by a doctoral research grant from the French Ministère de l’Enseignement Supérieur, de la Recherche et de l’Innovation (recipient: Maëlys Moulin), with additional support from Hydro-Québec (Canada) and EDF–RTE (France). The funders had no role in study design, data collection and analysis, decision to publish, or preparation of the manuscript.

## Ethical statement

The study received approval from the Institutional Review Board of the EuroMov Digital Health in Motion (DHM) Center (IRB-EM, approval no. 2111B) and was conducted in accordance with the Declaration of Helsinki. Written informed consent was obtained from all participants.

## Author contributions

M.M.: Conceptualization, Investigation, Methodology, Software, Formal analysis, Data curation, Visualization, Writing – original draft. E.F.: Methodology, Investigation. J.M.: Methodology, Validation. N.B.: Conceptualization, Supervision, Resources. S.R.: Conceptualization, Methodology, Software, Validation, Supervision. All authors: Writing – review & editing.

## Data availability

The datasets generated and analyzed during the current study are available from the corresponding author on request, to any qualified researcher.

## Declaration of generative AI technologies

During the preparation of this work, the authors used ChatGPT-5 (OpenAI, accessed January 2026) to assist in editing and rephrasing selected sections of the manuscript (Introduction and Discussion). In addition, ChatGPT-5.2 was used to generate the initial EEG icon used in Figure 2. The authors subsequently duplicated, edited, and recolored this icon and created the full figure layout. The authors reviewed and edited the content as needed and take full responsibility for the final version of the manuscript and figures.

## Supplementary data

Figure 5 details the subject-level quality-control procedure used to identify and exclude globally contaminated recordings.

**Figure 5.**
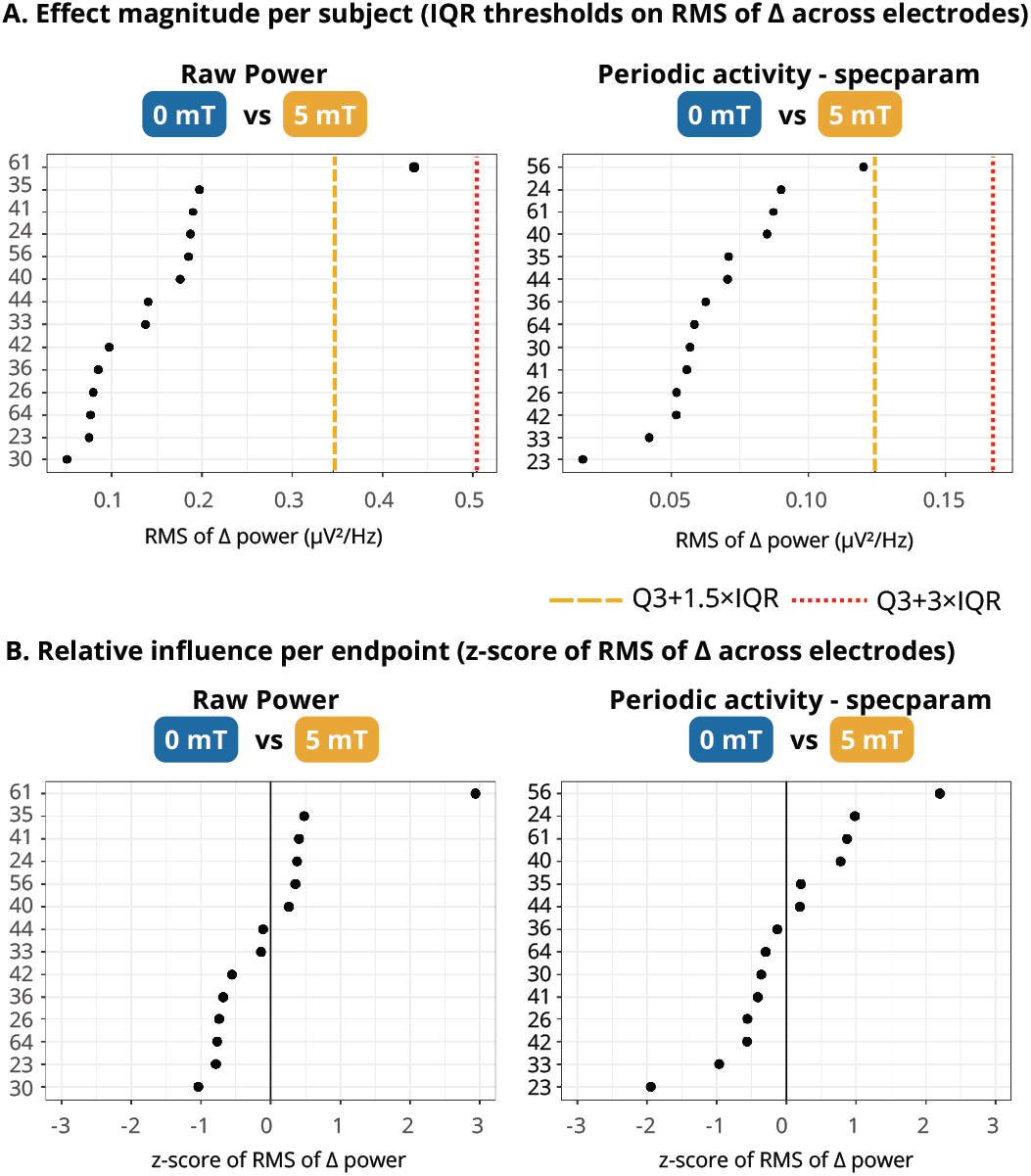
Data quality control and subject exclusion procedure. (A) Effect magnitude per subject, quantified as the root mean square (RMS) of the condition-related difference (Δ = 5 mT− 0 mT) across electrodes, for three representative EEG outcome measures: raw power and aperiodic-adjusted gamma power estimated using the *specparam* approach. Each dot represents one participant. Dashed orange and red vertical lines indicate outlier thresholds defined as *Q*_3_ + 1.5 *×*IQR and *Q*_3_ + 3 *×*IQR, respectively. (B) Relative influence of each participant, expressed as a *z*-score of the RMS of Δ across electrodes for the same EEG outcome measures. Positive *z*-scores indicate participants exerting a stronger-than-average influence on the group-level effect, whereas negative values indicate weaker contributions. One participant exceeded the predefined exclusion criterion and was excluded from subsequent analyses. Numbers on the vertical axis indicate participant identifiers. Participants were ordered independently within each panel according to the magnitude of the corresponding RMS effect, which explains why participant order may differ between panels.

Figure 6 shows the distribution of *specparam* goodness-of-fit (*R*^2^) values across stimulation conditions, providing a quality-control assessment of spectral parameterization performance.

**Figure 6.**
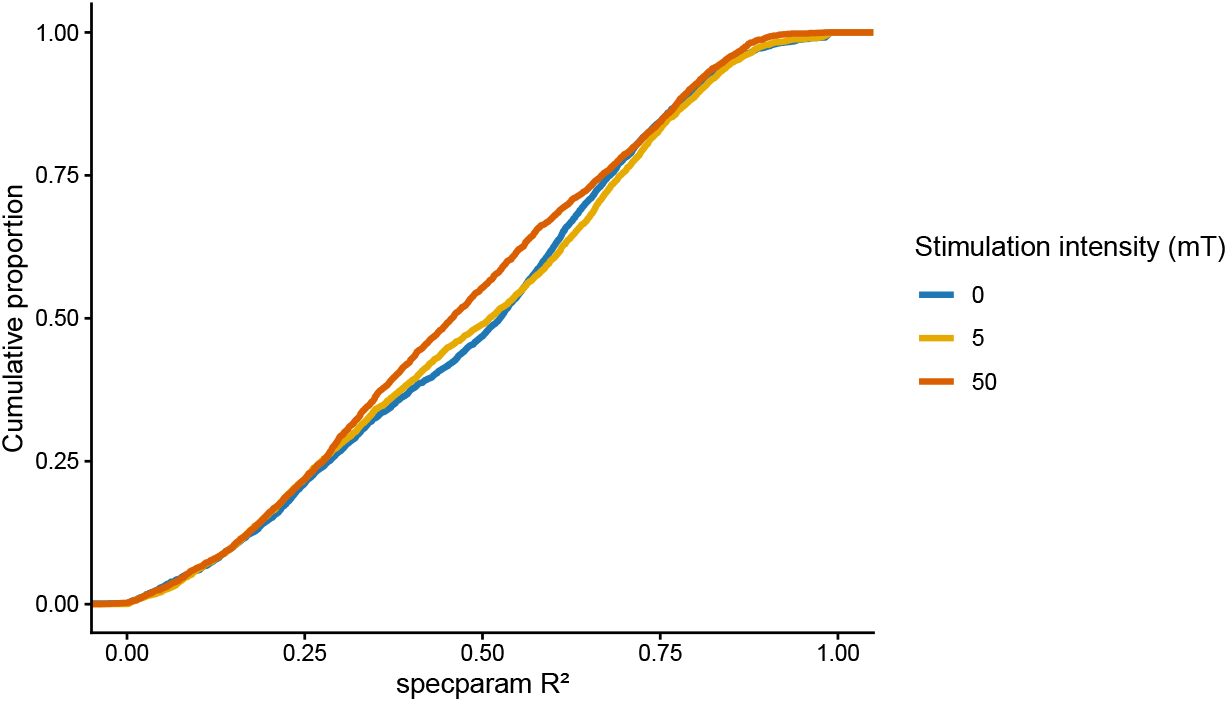
Distribution of spectral parameterization goodness-of-fit. Empirical cumulative distribution functions (ECDFs) of *specparam* goodness-of-fit (*R*^2^) values across stimulation intensities (0, 5, and 50 mT). The distributions show variable goodness-of-fit, with some low *R*^2^ values. This is expected because *specparam* models spectra as an aperiodic component plus Gaussian-like periodic peaks, whereas the present gamma-band effects were broadband and did not exhibit stable narrowband peaks. Fit quality may also be constrained by the restricted 30–80 Hz fitting window and by the exclusion of narrow frequency bands around 40, 50, and 60 Hz. Importantly, *R*^2^ distributions were broadly comparable across stimulation conditions, suggesting that condition-dependent differences in fit quality are unlikely to account for the observed effects. Accordingly, *specparam*-derived measures were interpreted as complementary aperiodic-adjusted gamma power estimates, and all main findings were compared with raw gamma power and log–log aperiodic-adjusted gamma power.

Figure 7 presents the full electrode-wise topographic comparison across stimulation contrasts and spectral representations, including effect sizes and Bonferroni-corrected statistics.

**Figure 7.**
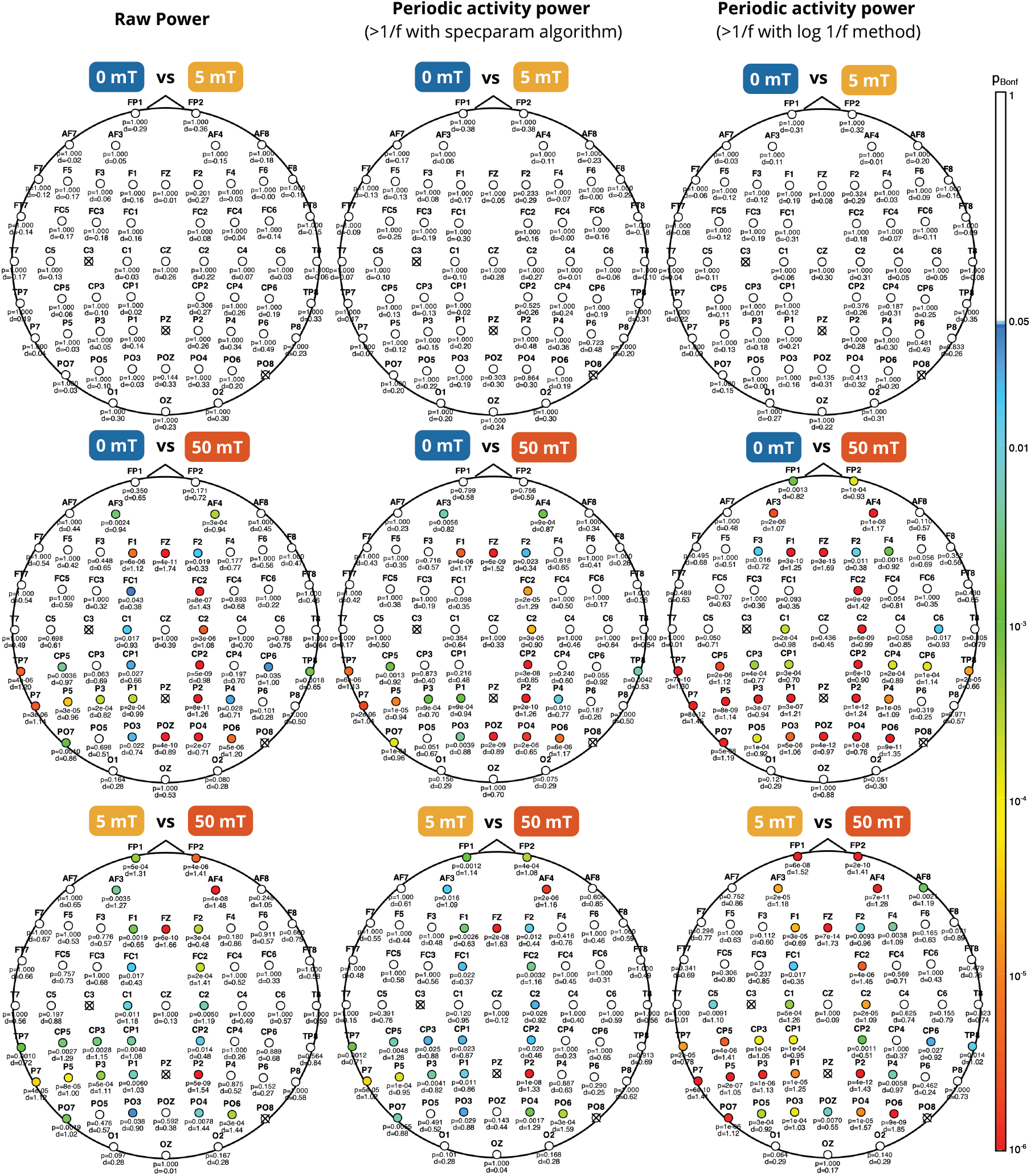
Scalp topographies of gamma-band power differences across stimulation intensities and spectral representations. Rows show within-subject contrasts (5 mT vs. 0 mT, 50 mT vs. 0 mT, and 50 mT vs. 5 mT), and columns correspond to raw gamma power (left), *specparam*-adjusted gamma power (middle), and log–log adjusted gamma power (right). Colored markers indicate electrodes reaching the significance threshold, with color encoding Bonferroni-corrected *p*-values on a log scale. Electrodes with corrected *p*≥ 0.05 are shown in white. Electrode-wise statistics are overlaid: *p* denotes the Bonferroni-corrected value and *d*_*z*_ the paired Cohen’s effect size. Excluded electrodes are shown as crossed markers.

## Notes

### Competing Interest Statement

The authors have declared no competing interest.

### Summary of Updates

This revised version includes several changes made to improve clarity, methodological transparency, and interpretation of the findings. We clarified the rationale for focusing the EEG analyses on three predefined stimulation conditions: 0 mT as the no-field control condition, 5 mT as a weak subthreshold exposure, and 50 mT as the robust suprathreshold condition. We also clarified that the 50 mT versus 0 mT contrast is the primary comparison, whereas the 50 mT versus 5 mT contrast is used as a secondary control comparison. We added a more explicit discussion of the limitation that the 50 mT condition combines high stimulation intensity with high perception frequency. The results are now interpreted as EEG correlates of a suprathreshold stimulation-perception state rather than as isolated perception-specific markers. We clarified the interpretation of specparam goodness-of-fit values. The revised manuscript explains that lower R2 values are expected because the observed gamma-band effects were broadband and did not show stable narrowband peaks, whereas specparam models spectra using an aperiodic component and Gaussian-like periodic peaks. We also emphasize that the main findings were compared across raw gamma power, specparam-adjusted gamma power, and log-log aperiodic-adjusted gamma power. We added clarifications regarding the interstimulus interval, the absence of a dedicated washout manipulation, and the pseudo-randomization of stimulation intensities. We also clarified that the absence of a focal occipital scalp effect does not rule out early visual cortex involvement, but indicates that such involvement, if present, was not expressed as a stable focal power change detectable at the scalp level. Finally, we shortened and reorganized the Discussion to reduce repetition, harmonized terminology throughout the manuscript, clarified figure captions, and proofread the references for consistency.

